# Faa1 membrane binding drives positive feedback in autophagosome biogenesis via fatty acid activation

**DOI:** 10.1101/2023.09.09.556965

**Authors:** Verena Baumann, Sonja Achleitner, Susanna Tulli, Martina Schuschnig, Lara Klune, Sascha Martens

**Author notes:** these authors contributed equally (Verena Baumann, Sonja Achleitner, Susanna Tulli).

## Abstract

Autophagy serves as a stress response pathway by mediating the degradation of cellular material within lysosomes. In autophagy this material is encapsulated in double membrane vesicles termed autophagosomes, which form from precursors referred to as phagophores. Phagophores grow by lipid influx from the endoplasmic reticulum into Atg9-positive compartments and local lipid synthesis provides lipids for their expansion. How phagophore nucleation and expansion are coordinated with lipid synthesis is unclear. Here, we show that Faa1, an enzyme activating fatty acids, is directly recruited to Atg9 vesicles. We further show that Faa1 binds to negatively charged membranes. We define the membrane binding surface in Faa1 and show that membrane binding is required for its enzymatic activity. In cells, membrane binding by Faa1 is required for its recruitment to phagophores and promotes autophagosome biogenesis. Our results suggest a positive feedback loop coupling phagophore nucleation and expansion to lipid synthesis.

**Summary:** Baumann, Achleitner, Tulli et al. dissect Faa1 function and recruitment during autophagy. They discover that Faa1 directly binds membranes via a positively charged surface. This is a prerequisite for Faa1’s enzymatic activity sustaining autophagosome biogenesis.

## Introduction

Macroautophagy (hereafter autophagy) mediates the degradation of harmful material within the lysosomal system. It also protects cells during times of starvation by the recycling of cellular material in order to provide energy and building blocks such as amino acids for the synthesis of essential factors. Autophagy therefore represents a major pillar of the cellular stress response and various diseases are associated with defects in this process[1, 2]. The hallmark of autophagy is the *de novo* formation of a double membrane organelle, the autophagosome. Upon induction of autophagy, the nucleation of a small membrane precursor referred to as phagophore (or isolation membrane) is initiated. This membrane structure gradually captures cytoplasmic material as it grows. After closure of the phagophore, the resulting double membrane autophagosome fuses with lysosomes (or the vacuole in yeast and plants) where the inner membrane and the cargo are degraded[3, 4].

The biogenesis of autophagosomes is mediated by a set of conserved factors, which can be grouped into the following modules (referring to *Saccharomyces cerevisiae* nomenclature): (I) the Atg1 kinase complex, (II) vesicles containing the Atg9 lipid scramblase, (III) the class III phosphatidylinositol 3-kinase complex 1 (PI3KC3C1) generating phosphatidylinositol 3-phosphate (PI3P), (IV) the Atg2 lipid transfer protein, (V) the PI3P binding PROPPINs Atg18 and Atg21 and (VI) the Atg8 lipidation machinery including the Atg12–Atg5-Atg16 complex, which mediates the attachment of the ubiquitin-like Atg8 protein to the headgroup of phosphatidylethanolamine (PE)[5–8]. Over the past years a consensus model for how this machinery and its biochemical activities act in concert to mediate the formation and expansion of phagophores has emerged. According to this model phagophore formation is initiated in proximity to the endoplasmic reticulum (ER) by Atg9 containing membrane precursors[9–17]. The ER and the Atg9 positive membrane precursors are connected by the Atg2 lipid transfer protein[13, 15, 18, 19]. The Atg1 kinase complex and the PI3KC3C1 in conjunction with the PROPPINs facilitate the formation of these membrane contact sites and promote the subsequent expansion of these initial membrane seeds by recruiting and activating the Atg8 lipidation machinery[20, 21]. This machinery in turn mediates the conjugation of Atg8 to PE on the expanding phagophore[22–34].

The main lipid source for autophagosome biogenesis is the ER from where the lipids are transported to the phagophore by the Atg2 lipid transfer protein[35–38]. Another lipid transfer protein, Vps13, is partially redundant with Atg2 with respect to the lipid transfer activity[39]. In the phagophore, the ER derived lipids are distributed between the two leaflets by the Atg9 lipid scramblase[40–42]. It has been shown that lipid synthesis in the ER is required to provide sufficient lipids for autophagosome biogenesis[43–45]. In particular, it has been demonstrated both in yeast and in mammalian cells that *de novo* synthesized lipids in the ER can be detected in considerable amounts in autophagosomal membranes[43, 44, 46]. In addition, in mammalian cells autophagosome biogenesis takes place at ER sites, which are enriched in phosphatidylinositol (PI) synthase (PIS)[47]. In *S. cerevisiae* the long-chain fatty acid-coenzyme A (acyl-CoA) synthetases (ACSs) Faa1 and Faa4, which catalyze the first step in lipid synthesis by generating acyl-CoA, localize to phagophores. Their catalytic activities are redundant with each other and with the fatty acid synthase (FAS). Upon inhibition of FAS activity either by glucose starvation or cerulenin treatment, the recruitment of Faa1 as well as its catalytic activity becomes critical for autophagosome biogenesis. It has been shown that the recruitment of Faa1 to Atg8 positive structures at detectable levels depends on Atg1, Atg5, Atg9 and Atg14[43]. Yet, the mechanistic details of Faa1 recruitment and activation on membranes remain elusive.

Here, we set out to characterize the membrane recruitment mechanism of Faa1 and its relevance for autophagosome formation. We found that Faa1 binds to Atg9 vesicles isolated from yeast as well as artificially generated proteoliposomes containing Atg9. Furthermore, Faa1 is directly recruited to negatively charged lipids such as PI as well as PI3P and its catalytic activity relies on this membrane binding. Mutations in the membrane binding site abolish the interaction of Faa1 with membranes *in vivo*, decrease autophagic flux and reduce cell survival upon starvation and FAS inhibition. Our results suggest a positive feedback loop where the recruitment of Faa1 to Atg9 vesicles and phagophores activates fatty acids for local lipid synthesis to sustain phagophore expansion. In turn, phagophore expansion creates further binding sites for Faa1.

## Results

### Faa1 is recruited to Atg9 vesicles and directly binds negatively charged membranes

Faa1 accumulates on phagophores where it is proposed to locally activate fatty acids for lipid synthesis in the ER. The generated phospholipids are subsequently transported into the expanding phagophore by Atg2 (Figure 1A)[43]. The membrane recruitment and activation of Faa1 is thus a crucial step in autophagosome biogenesis. Interestingly, proteomic analyses have identified Faa1 in Atg9 vesicle enriched fractions [9], suggesting an early role for Faa1 during autophagosome biogenesis. In order to obtain insights into the mechanistic basis of Faa1 recruitment to the site of autophagosome biogenesis, we tested if Faa1 could directly bind to Atg9 vesicles. To this end, we isolated Atg9 vesicles from yeast cells and tethered them via EGFP-tagged Atg9 to GFP-trap beads. Upon incubation with Faa1-mCherry purified from *S. cerevisiae*, we observed binding of Faa1 to the vesicle-covered beads by confocal microscopy (Figure 1B). Atg9 vesicles have been shown to act as an assembly platform for the downstream autophagy machinery. Among the tethered factors are lipid binding proteins like Atg21 and Atg18, that directly bind PI3P generated by the PI3KC3C1. Therefore, we tested whether the recruitment of Faa1 could be enhanced by an active PI3KC3C1. However, comparison of Faa1 binding in presence of ATP/AMP-PNP and PI3KC3C1 revealed no difference in this setup.

**Fig. 1.**
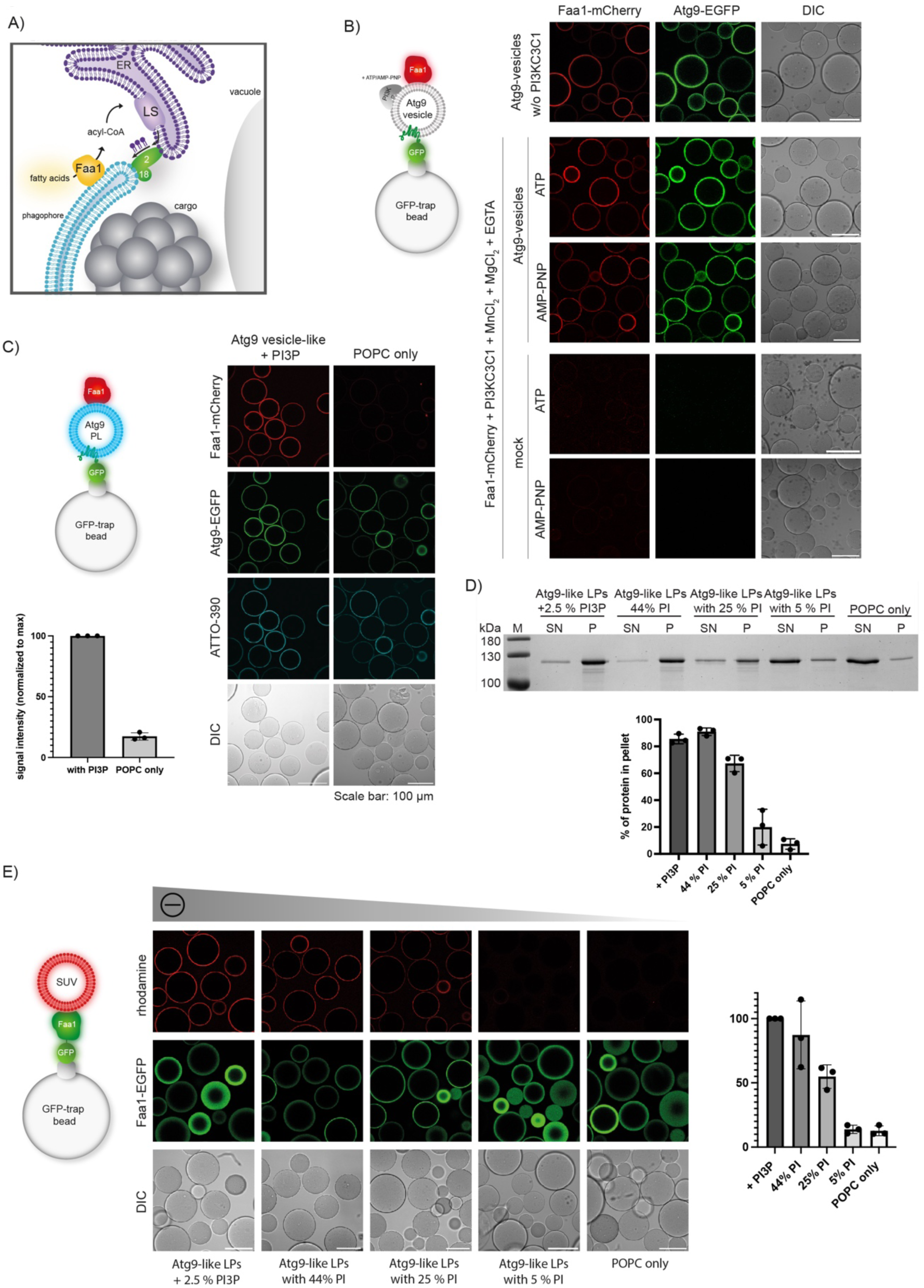
Faa1 directly binds negatively charged membranes. (A) Model for autophagosome expansion driven by localized de novo phospholipid synthesis. (B) Faa1 is recruited to Atg9 vesicles isolated from cells independent of PI3KC3C1. The experimental setup is shown on the left. (C) Faa1 can be recruited to Atg9 PLs. GFP-Trap beads were coated with Atg9-EGFP PLs and incubated with 1 µM Faa1-mCherry as depicted on the left. Quantification of Faa1 recruitment to Atg9 PLs can be seen on the bottom left (n=3). (D) Co-sedimentation assay showing that Faa1 is recruitment to Atg9 vesicle-like liposomes dependent on the net charge of the membranes (n=3). (E) Setup of the microscopy-based membrane recruitment assay: Faa1-EGFP is immobilized on GFP-trap beads, incubated with 1 mM of rhodamine labelled liposomes and imaged by microscopy. In accordance with the results in 1D, the degree of liposome (LPs/SUVs) recruitement to the Faa1-EGFP-covered beads depends on the net charge of the membranes (n=3). Liposome composition for 1D and 1E resembles Atg9 vesicles (44% POPC, 6% POPS, 44% liver PI, 5.5% POPE, 0.5% lissamine rhodamine-PE). For the liposomes containing PI3P, the percentage of PI was reduced while a decreased percentage of PI was substituted for POPC. The quantifications of 1C, 1D and 1E depict the mean and standard deviation.

Yeast Atg9 vesicles have a very distinct lipid composition characterized by a high PI content (44%)[9]. Aiming to distinguish if Faa1 recruitment to Atg9 vesicles occurs through direct membrane binding or interactions with other proteins, we generated artificial proteoliposomes (PLs). These PLs mirror the native lipid composition of Atg9 vesicles + PI3P and contain Atg9 but lack additional proteins. As a control PLs only consisting of POPC and Atg9-EGFP were used. Atg9-PLs were immobilized on GFP-trap beads and incubated with Faa1-mCherry. The liposome membrane was labelled in blue by addition of ATTO-390-DOPE (Figure 1C). While we observed robust Faa1 recruitment to PL with an Atg9 vesicle-like lipid composition, POPC only Atg9-PLs showed barely any binding of Faa1. This indicates that Faa1 membrane binding is mainly driven by its direct interaction with specific phospholipids.

To determine which lipids are bound by Faa1 we first employed a liposome co-sedimentation assay (Figure 1D). The used liposomes mimicked an Atg9 vesicle lipid composition with and without PI3P. Both liposome mixes robustly bound Faa1 independent of other proteins. Systematic reduction of negatively charged PI resulted in a concurrent decline of membrane recruitment. Faa1 tethering to liposomes comprised solely of POPC was barely detectable. To corroborate our findings, we conducted a microscopy-based membrane protein interaction assay. In this experimental setup, we immobilized Faa1-EGFP on GFP-trap beads and incubated them with the liposome compositions described in Figure 1, supplemented with lissamine rhodamine-PE to visualize the liposomes. Consistent with the co-sedimentation assay, we observed an increased Faa1 recruitment associated with higher levels of negative charge in the liposomes. In conclusion, our results suggest that Faa1’s membrane recruitment is primarily driven by negatively charged lipids including PI and PI3P.

### A positively charged surface on Faa1 mediates membrane binding

The results above indicated that Faa1 directly binds negatively charged lipids. This led us to hypothesize that a positively charged, surface-exposed region of Faa1 is responsible for the protein-lipid interaction. Since no experimental structure is available for the Faa1 protein or any of its close homologs, we made use of the structure predicted by AlphaFold2 (https://alphafold.ebi.ac.uk/entry/P30624) to identify the putative membrane binding site. The calculated electrostatic surface potential indeed suggested the presence of a positively charged surface on the protein (Figure 2A). Interestingly, this surface is also present in human FACL4, the closest relative to Faa1 in humans (Figure S1). In order to test if this region is responsible for membrane binding, we mutated lysine residues located in the corresponding surface patch to aspartate. In particular, we generated a 4D mutant with K387, K388, K636 and K647 and a 6D mutant with K612 and K619 additionally mutated to aspartate. The corresponding mutant proteins were expressed and purified. The 4D and 6D mutants behaved identical to the wild type protein during the purification including size exclusion chromatography, where all proteins eluted as monomers suggesting that the mutations did not result in protein misfolding and aggregation (Figure S2). We then tested the mutants for their ability to bind to membranes in a liposome co-pelletation assay (Figure 2B). As expected, the wild type protein robustly co-pelleted with liposomes containing an Atg9 vesicle-like lipid composition including PI3P. Both mutants showed a significantly reduced co-sedimentation with these liposomes (Figure 2B), suggesting that the mutated surface is a major contributor to membrane binding. We corroborated these results with a microscopy-based membrane binding assay (Figure 2C), where the 4D and 6D mutants also showed a significantly reduced binding activity towards Atg9 vesicle-like liposomes. We conclude that a conserved positively charged surface on Faa1 mediates membrane binding to negatively charged membranes.

**Fig. 2.**
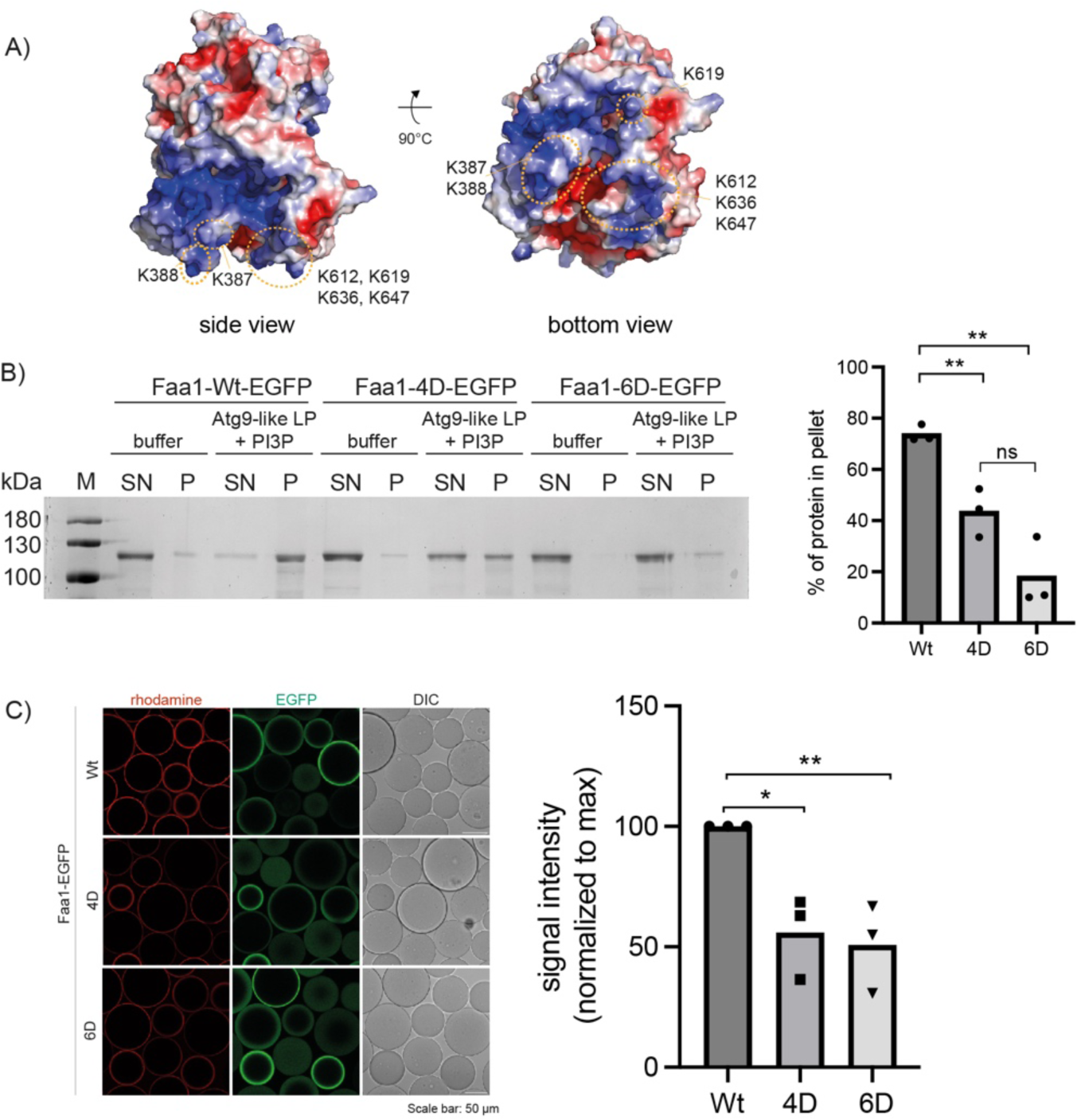
A positively charged surface area mediates Faa1 membrane binding in vitro. (A) Alphafold model of scFaa1. Electropositively and electronegatively charged areas are colored in blue and red, respectively. Neutral residues are in white. Mutated amino acids are highlighted with orange circles. (B) Co-sedimentation assay showing that mutations of positively charged residues in the plane area (Faa1-4D, Faa1-6D) significantly reduce membrane binding compared to Faa1-Wt (n=3). (C) Microscopy-based membrane recruitment assay with different versions of Faa1. Faa1 is immobilized on GFP-trap beads, incubated with 1 mM of rhodamine labelled liposomes and imaged by microscopy (n=3). Liposome composition for 2B and 2C resembles Atg9 vesicles + PI3P (44% POPC, 6% POPS, 41.5% liver PI, 5.5% POPE, 2.5% PI3P + 0.5% lissamine rhodamine-PE). The quantifications depict the mean and standard deviations. t-test: *<0.05, **<0,01.

### The enzymatic activity of Faa1 requires membrane binding

Next, we asked whether the interaction of Faa1 with membranes affects its enzymatic activity. To conveniently follow the activity of Faa1, we employed a coupled enzymatic assay[48]. This assay uses the AMP that is produced by Faa1 during the ATP-dependent acyl-CoA production for the oxidation of NADH to NAD^+^. In contrast to NAD^+^, NADH absorbs at 340 nm and therefore the conversion of NADH to NAD^+^ can be followed spectroscopically (Figure 3A). In the absence of Faa1 almost no reduction in absorption at 340 nm was detected, while the positive control with directly added AMP resulted in very fast reduction of absorption (Figure 3B). Upon addition of Faa1 and POPC liposomes the absorbance was comparable to the negative control. However, the addition of liposomes with an Atg9 vesicle-like lipid composition with and without PI3P did promote Faa1 activity as measured by a decreased absorbance at 340 nm over time. This suggests that membrane binding by Faa1 is required for its activity. Notably, the requirement of membrane binding for the enzymatic activity of Faa1 is not a general property of ACSs because FadD, a bacterial ACS derived from *E. coli* is active in the absence of liposomes (Figure 3C). We then went on to test if the identified positively charged surface patch is required for the membrane-induced activation of the enzymatic activity of Faa1. Indeed, wild-type Faa1 consistently exhibited enzymatic activity, whereas both mutants displayed no detectable activity beyond the negative control (Figure 3D), highlighting the significance of the identified region in mediating not only membrane interaction but also Faa1 activity.

**Fig. 3.**
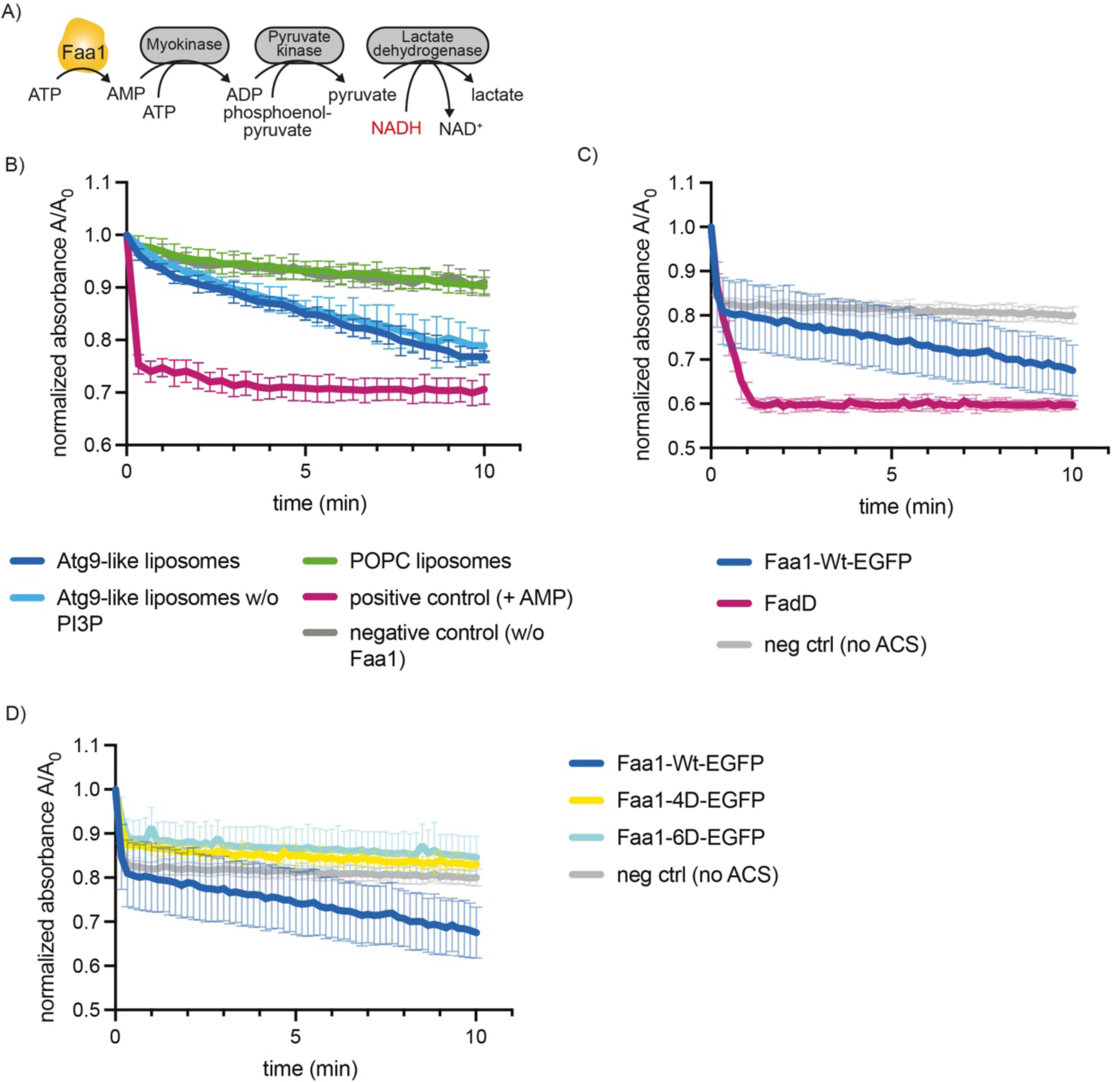
Membrane binding of Faa1 is important for its activity. (A) Scheme of the coupled enzymatic assay for testing the activity of Faa1. (B) Coupled enzymatic assay following the activity of Faa1 in the presence of different liposomes by measuring absorbance of NADH at 340 nm. (C) Coupled enzymatic assay comparing the activities of FadD in the absence of liposomes and Faa1 in the presence of Atg9 vesicle-like liposomes. (D) Enzymatic assay comparing Faa1 to Faa1-4D and Faa1-6D. Liposome composition for 3C-3D resemble Atg9 vesicles + PI3P (44% POPC, 6% POPS, 41.5% liver PI, 6% POPE, 2.5% PI3P).

### Faa1 membrane binding is crucial for its localization to autophagosomes, cell survival and autophagic flux

After discovering the crucial role of the positively charged surface area of Faa1 in membrane binding and fatty acid activation, our focus shifted to understanding the implications of these findings *in vivo*. In yeast cells, Faa1 had previously been observed to localize to various cellular compartments, including the plasma membrane, ER and autophagosomes[39]. To investigate the impact of mutating the identified positively charged region on the localization of Faa1, we employed genomically modified yeast strains. These strains co-expressed different variants of Faa1-3xGFP along with the mCherry-tagged autophagy marker, Atg8. In line with previous observations, our live-cell imaging showed wild type Faa1 being recruited to the plasma membrane[43], the ER and Atg8-positive autophagosomal membranes upon autophagy induction with rapamycin. In contrast, both Faa1-4D and Faa1-6D did not bind membranes but remained cytosolic (Figure 4A). Importantly, the expression levels of these mutants were comparable to those of the wild-type protein (Figure S3A).

**Fig. 4.**
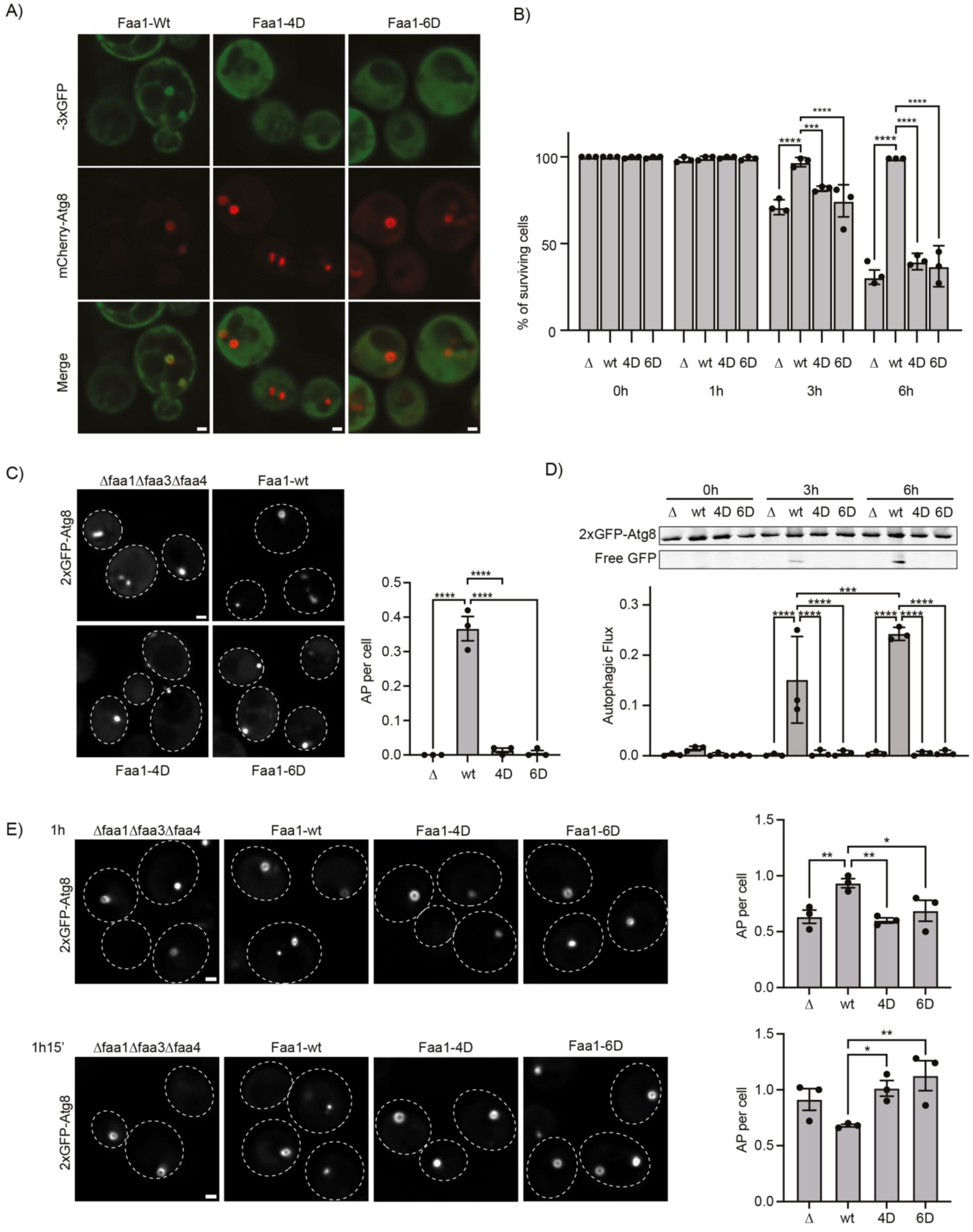
Mutation of positively charged residues in membrane binding surface reduces Faa1 membrane localization and function in vivo. (A) Faa1, Faa1-4D and Faa1-6D localization in cells co-expressing mCherry-ATG8. Indicated strains were imaged via fluorescence microscopy after rapamycin treatment (1 h). n=3, scale bars, 1µm. (B) Survival of indicated strains during starvation + cerulenin 0, 1, 3, and 6h of treatment. Data are means ± SD (n = 3; 100 cells/strain per replicate). (C) Fluorescent microscopy assessment of autophagosome biogenesis in indicated strains after 1h nitrogen starvation + cerulenin. Left, representative images of the analyzed strains. Dashed lines indicate cell boundaries. Scale bar, 1µm. Right, respective quantification of autophagosomes (AP). Data are means ± SEM (n = 3; 50 cells/strain per replicate). (D) Autophagic flux of indicated strains expressing 2xGFP-ATG8 after 0, 3, 6h of starvation + cerulenin. Data are means ± SD (n = 3). (E) Fluorescent microscopy assessment of autophagosome biogenesis in indicated strains after 1h and 1h15’ of nitrogen starvation and glucose depletion (0.01% w/v). Left, representative images of the analyzed strains at indicated time points. Dashed lines indicate cell boundaries. Scale bar, 1µm. Right, respective quantification of AP. Data are means ± SEM (n = 3; 50 cells/strain per replicate). One-way ANOVA: *<0.05, **<0.01, ***<0.001.

We subsequently investigated, the potential impact of the membrane-binding mutants on autophagy. To this end, we co-expressed 2xGFP-Atg8 alongside either wild type Faa1 or the mutant variants in starved *faa1Δfaa3Δfaa4Δ* cells. Autophagic flux was monitored utilizing the GFP-Atg8 assay, which exploits the distinct stabilities of Atg8 and GFP within the vacuole[49]. Upon fusion of the autophagosome with the vacuole to form an autolysosome, Atg8 is rapidly degraded while GFP remains intact for a longer time and can be detected as free GFP by western blot analysis. Under these conditions, we did not observe any difference in the autophagic flux between Faa1 wild type or mutants (Figure S3B). Notably, in yeast cells, the activation of fatty acids can be carried out by both, ACSs and FAS via two parallel pathways [50]. Furthermore, ACSs activity is key for the survival of starving cells upon FAS inhibition via cerulenin. We therefore examined whether Faa1-4D and Faa1-6D could restore the viability of starved and cerulenin treated *faa1Δfaa3Δfaa4Δ* cells. As expected, wild type Faa1 expression fully rescued cell survival. However, cells expressing Faa1-4D or Faa1-6D were unable to sustain cell viability under these conditions (Figure 4B).

We then proceeded to evaluate the effect of the membrane binding mutants on autophagosome biogenesis and autophagic flux upon FAS inhibition. Autophagosome formation was monitored by live cell imaging of 2xGFP-Atg8 (Figure 4C). Cells expressing Faa1-4D or Faa1-6D hardly showed any autophagosomes, suggesting severely impaired autophagy. To further corroborate these findings, we monitored autophagic flux by employing the GFP-Atg8 assay. As expected, we observed efficient GFP-Atg8 cleavage in cells expressing wild type Faa1 (Figure 4D). Conversely, no free GFP could be detected in cells expressing Faa1-4D or Faa1-6D.

As an alternative and physiologically relevant way to inhibit FAS, we lowered the glucose concentration (0.01%) during starvation and assessed Faa1-4D and Faa1-6D function (Figure 4E). Under this metabolic condition, the effect of the Faa1-4D and Faa1-6D mutations was less prominent compared to the response seen with cerulenin treatment. However, when cells were imaged to assess autophagosome biogenesis after 1 h of starvation in reduced glucose, Faa1-4D and Faa1-6D cells showed fewer autophagosome than wild type Faa1. Interestingly, a reversal in this trend was observed after 1 h and 15’ with Faa1-4D and Faa1-6D cells showing more autophagosomes than wild type Faa1 cells (Figure 4E). This suggests that impaired Faa1 function, combined with reduced glucose levels during starvation results in slower autophagosome biogenesis. Since reduction of glucose during starvation did not entirely suppress autophagosome formation, we proceeded to assess autophagic flux. Our analysis revealed a reduction compared to wild type Faa1 after 3 hours when Faa1-4D and Faa1-6D were expressed (Figure S3C).

Taken together, these findings show that the identified positively charged surface region on Faa1 plays a pivotal role in facilitating its binding to membranes *in vivo*. Modification of the respective residues renders Faa1 cytosolic and comprises the cell’s capacity to sustain cell survival under starvation and FAS inhibition, as well as autophagic flux.

## Discussion

ACSs, including the long-chain fatty acid-CoA synthetase Faa1, kick-start phospholipid synthesis by catalyzing the formation of acyl-CoA from long-chain fatty acids[50]. Besides its localization to the plasma membrane and the ER, Faa1 was also detected on phagophores. Re-localization of Faa1 from the site of autophagosome biogenesis to the plasma membrane results in an impairment of autophagic flux, underlining the pivotal role of localized *de novo* lipid synthesis for phagophore expansion[43].

Here, we sought to elucidate the mechanisms behind Faa1 recruitment to cellular membranes and its activation. Our findings revealed that Faa1 binds Atg9 vesicles isolated from cells. These vesicles do not only contain Atg9 but several other components including Atg23 and Atg27[9]. To determine the factors necessary and sufficient for Faa1 recruitment, we therefore made use of more controlled and simplified setups. We employed Atg9-PL, which contain only phospholipids and Atg9 but are deprived of other proteins. In this experimental set up Faa1 was still recruited to Atg9 vesicle-like PL although in an Atg9 independent manner. This indicates that membrane composition rather than Atg9 plays a crucial role for the localization of Faa1. Indeed, experiments testing Faa1’s binding to liposomes showed that the protein has a clear preference for membranes containing negatively charged phospholipids such as PI and PI3P.

In the cell, organelles exhibit highly distinctive membrane compositions[51]. These compositions play a pivotal role in governing the spatiotemporal control of cellular processes. [52, 53]. PI3P for example is primarily found on endosomes and autophagic membranes, assuming a central role in the endocytic pathway by orchestrating endosomal fusion. Atg9 vesicles have been shown to be particularly rich in PI[9]. This does not only make them a good substrate for the PI3KC1C3 but also a suitable recruitment platform for Faa1. Direct membrane-protein interactions like this are generally based on electrostatic forces between negatively charged lipids and positively charged regions on the protein surface [52]. By employing both, in vitro reconstitution assays as well as cell biological experiments, we identified such a positively charged surface on Faa1 as the mediator of its membrane binding activity. Mutation of the positive residues to aspartate (Faa1-4D and Faa1-6D) results in a loss of Faa1 recruitment to negatively charged membranes. The relatively flat, positively charged surface patch on Faa1 is consistent with a rather non-specific preference for the binding to negatively charged phospholipids. Interestingly, an analysis of the AlphaFold prediction of FACL4, the closest mammalian homologue of Faa1, revealed that the positively charged surface region is conserved. This suggests a similar membrane recruitment mechanism of FACL4 in mammalian cells.

Our data further revealed that in addition to correct localization, direct membrane binding by Faa1 is also a prerequisite for its enzymatic activity. This is in contrast to the bacterial homologue of Faa1, FadD, which does not require any membranes to be active. An impairment of Faa1’s interaction with membranes in vivo resulted in reduced autophagic flux as well as compromised autophagosome formation and reduced cell survival during conditions of starvation and FAS-inhibition. The structural and mechanistic basis for the membrane binding-dependent enzymatic activity of Faa1 remains to be determined.

The dependence of membrane binding for the activation of Faa1 offers another layer of regulation to initiate lipid synthesis at the right time and space. In autophagy, this dependence allows positive feedback where the initial recruitment of Faa1 to Atg9-positive phagophore seeds initiates lipid synthesis (Figure 5A-C). The synthesized lipids support membrane expansion creating additional binding sites for Faa1 and thus more products for lipid synthesis. Faa1 mutants that are unable to bind the membrane fail to fuel this feedback loop. The ability of Faa1 to directly bind negatively charged membranes further suggests that its targeted recruitment to different membranes including endosomes or the phagophore requires an additional level of regulation such as an interaction partner in order to prevent uncoordinated lipid synthesis within the cell. Regardless of the potential supplementary targeting factor, the stable membrane anchoring of Faa1 relies on the interaction between the positively charged surface region and negatively charged phospholipids.

**Fig. 5.**
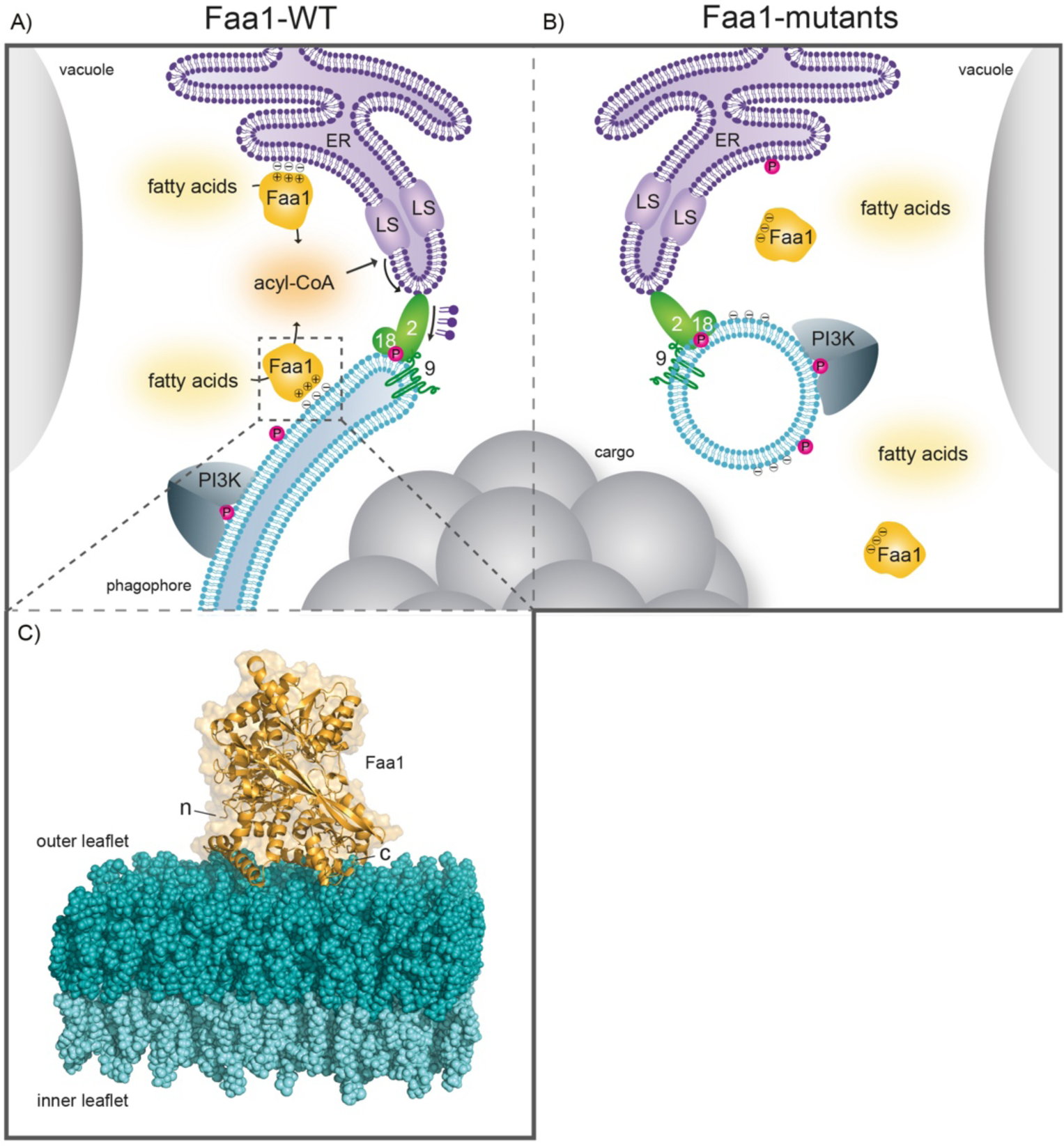
Model for the role of Faa1 during autophagosome formation and the consequences of its impaired membrane binding. (A) Faa1 is recruited to membranes upon presence of negatively charged lipids. There, it locally activates free fatty acids. The generated acyl-CoA enters the lipid synthesis machinery at the ER where it is converted to different types of phospholipids. Atg2 transfers the produced phospholipids into the growing phagophore where their distribution between outer and inner leaflet is re-established via the Atg9 lipid scramblase. The growing membrane provides further binding sites for Faa1. (B) Faa1-4D and Faa1-6D mutants fail to localize to membranes due to mutations in the positively charged surface area. This leads to a loss of Faa1 activity and further an impairment of autophagic flux and cell survival upon cerulenin treatment. (C) Model of Faa1 membrane interaction with Faa1 in yellow and the membrane bilayer in dark and light teal. Model was generated with CHARMM-GUI HMMM Builder[54, 55].

## Acknowledgements

We thank Martin Graef for critical discussions and support. We acknowledge funding by the Austrian Science Fund (FWF grants P35061-B, P32814-B). We thank the Max Perutz Labs BioOptics and Mass Spectrometry facility for technical support. Proteomics analyses were performed using the VBCF instrument pool.

## Author contributions

S.M. designed and supervised research, V.B., S.A., and S.T., designed research, V.B., S.A., S.T., L.K. and M.S. performed research. All authors analyzed data and commented on the manuscript. S.M. and V.B. wrote the manuscript.

## Materials and Methods

### Protein expression and purification

A list of constructs for protein expression can be found in Table 1. The purification procedures of proteins from constructs generated in this study are described below.

**Table 1.** Table of yeast strains.

| Name | Genotype | Background | Reference |
| --- | --- | --- | --- |
| BY4741 | MAT a; *hi3Δ1 leu2Δ0 met15Δ0 ura3Δ0* |  | Euroscarf |
| W303 | *ade2-1; leu2-3; his3-11, 15; trp1-1; ura3-1; can1-100* |  | Thomas and Rothstein (1989) |
| SMY1 |  | BY4741 | Euroscarf |
| SMY276 | MAT a; *pep4::NAT ATG9-EGFP-TAP:URA* | BY4741 | [9] |
| SMY462 | *faa1::KANMX4* | BY4741 | Euroscarf |
| SMY464 | *faa1::KANMX4 leu2::GAL1pr-FAA1-mCherry-TAP-ADH1ter:LEU2* | BY4741 | This study |
| SMY465 | *faa1::KANMX4 leu2::GAL1pr-FAA1-EGFP-TAP-ADH1ter:LEU2* | BY4741 | This study |
| SMY503 | *faa1::KANMX4 leu2::GAL1pr-faa1-K387D/K388D/K636D/K647D-EGFP-TAP-ADH1ter:LEU2* | BY4741 | This study |
| SMY506 | *faa1::KANMX4 leu2::GAL1pr-faa1-K387D/K388D/K612D/K619D/K636D/K647D-EGFP-TAP-ADH1ter:LEU2* | BY4741 | This study |
| SMY511 | *Δatg19::HIS3 pRS304-prATG8-mCherry-ATG8 FAA1-3xyEGFP-CaURA3* | W303 | Schütter et al. (2020)[43] |
| SMY512 | *Δatg19::HIS3 pRS304-prATG8-mCherry-ATG8 Faa1- K387D/K388D/K636D/K647D -3xyEGFP-CaURA3* | W303 | This study |
| SMY513 | *Δatg19::HIS3 pRS304-prATG8-mCherry-ATG8 Faa1- K387D/K388D/K612D/K619D/K636D/K647D -3xyEGFP-CaURA3* | W303 | This study |
| SMY514 | *Δatg19::natMX6 Δfaa1::FAA1-kanMX6 Δfaa3::TRP1 Δfaa4::HIS3 pRS306-prATG8-2xyEGFP-ATG8* | W303 | Schütter et al. (2020)[43] |
| SMY515 | *Δatg19::natMX6 Δfaa1::Faa1-K387D/K388D/K636D/K647D-kanMX6 Δfaa3::TRP1 Δfaa4::HIS3 pRS306-prATG8-2xyEGFP-ATG8* | W303 | This study |
| SMY516 | *Δatg19::natMX6 Δfaa1::Faa1-K387D/K388D/K612D/K619D/K636D/K647D -kanMX6 Δfaa3::TRP1 Δfaa4::HIS3 pRS306-prATG8-2xyEGFP-ATG8* | W303 | This study |
| SMY517 | *Δatg19::natMX6 Δfaa1::TRP1 Δfaa3::kanMX6 Δfaa4::HIS3 pRS306-prATG8-2xyEGFP-ATG8* | W303 | This study |

**Table 2.** Table of constructs.

| Name | Expressing | Vector | Expression system | Reference |
| --- | --- | --- | --- | --- |
| SMC1230 | 6xHis-TEV-Atg9-mEGFP2xStrep | pFastBac HT | Sf9 cells | Sawa-Makarska et al. (2020)[9] |
| SMC1496 | FadD-TEV-6xHis | pET-Duet | *E. coli* Rosetta pLysS | This study |

**Table 3.** Table of lipids.

|  | Name | Supplier | Catalog number | concentration |
| --- | --- | --- | --- | --- |
| POPC | 1-palmitoyl-2-oleoyl-glycero-3-phosphocholine | Avanti Polar Lipids, Inc. | 850457C | 10 mg/mL |
| POPS | 1-palmitoyl-2-oleoyl-sn-glycero-3-phospho-L-serine (sodium salt) | Avanti Polar Lipids, Inc. | 840034C | 10 mg/mL |
| POPE | 1-palmitoyl-2-oleoyl-sn-glycero-3-phosphoethanolamine | Avanti Polar Lipids, Inc. | 850757 | 10 mg/mL |
| Liver PI | L-α-phosphatidylinositol (Liver, Bovine) (sodium salt) | Avanti Polar Lipids, Inc. | 840042C | 10 mg/mL |
| PI3P | 1,2-dioleoyl-sn-glycero-3-phospho-(1'-myo-inositol-3'-phosphate) (ammonium salt) | Avanti Polar Lipids, Inc. | 850150P | 1 mg/mL |
| Rhodamine PE | Lissamine<sup>TM</sup> rhodamine B 1,2dihexadecanoyl-sn-glycero-3phosphoethanolamine, triethylammonium salt | Invitrogen | L-1392 | 1 mg/mL |
| ATTO 390 | ATTO390-labelled 1,2-Dioleoyl-sn-glycero3-phosphoethanolamine | ATTO-Tec | AD390-161 | 10 mg/mL |

Faa1-mCherry, Faa1-EGFP, Faa1-4D-EGFP and Faa1-6D-EGFP were purified from the SMY464, SMY465, SMY503 and SMY506 strains, respectively. Cells were grown at 30 °C in YPG up to an OD600 of 7-11. Cells were pelleted, washed once with cold H_2_O, once with lysis buffer A (500 mM NaCl, 50 mM Hepes pH 7.5, 2 mM MgCl_2_) and finally resuspended in lysis buffer containing complete protease inhibitors (Roche), an FY-inhibitor mix (Serva), DNAse I (Sigma), benzonase (Sigma) and 1 mM DTT. Resuspended cells were frozen in liquid nitrogen as pearls and lysed via freezer milling. The milled powder was thawed and resuspended in lysis buffer A by rolling gently at 4 °C. Lysates were cleared by centrifugation (20,000 rpm for 45 min at 4°C in a Beckman SW45Ti rotor). The supernatant was incubated with IgG Sepharose beads (GE Healthcare) on a rotary wheel for 1 h at 4 °C. Beads with bound protein were washed twice with lysis buffer A, once with high salt buffer (700 mM NaCl, 25 mM HEPES + DTT) and twice with lysis buffer A without protease inhibitors. Bound protein was cut off the beads with TEV protease for 1 h at 16 °C. The cleaved protein was concentrated and loaded onto a Superose 6 Increase size exclusion chromatography column (10/300 prep grade, GE Healthcare). Elution was carried out with 150 mM NaCl, 25 mM HEPES pH 7.5 and 1 mM DTT buffer. Fractions containing the protein of interest were pooled, concentrated, frozen in liquid nitrogen, and stored at −70°C.

Expression and purification of Atg9-EGFP and the PI3KC1C3 as well as generation of Atg9-PLs were carried out as described in Sawa-Makarska et al. 2020. Atg9-PL compositions were as follows: Atg9-vesicle like liposomes + PI3P (42.5 % POPC, 6 POPS, 6 % POPE, 41.5 % liver PI and 2.5 % PI3P and 1.5 % ATTO 390), POPC only liposomes (98.5 % POPC, 1.5 % ATTO 390).

### Atg9 vesicle isolation

Atg9 vesicle isolation was mainly carried out according to Sawa-Makarska et al. 2020 with minor optimization steps[9]: Briefly, binding of the cleared Atg9-EGFP vesicles to the IgG beads was carried out o/n. Cleavage from the beads was carried out for 3 hours at 4°C. The final reaction buffer was changed to 135 mM NaCl, 25 mM Tris.

For the microscopy-based interaction assay between Atg9 vesicles and Faa1, bead-bound Atg9 vesicles were incubated for 2 hours at RT with and without 50 nm of PI3KC1C3 including 0.5 mM MgCl2, 2 mM MnCl2, 1 mM EGTA and 0.2 mM ATP/AMP-PNP. Next, 1 µM of Faa1-mCherry was added and incubated for 2 hours. Imaging was carried out as described for the microscopy-based protein-membrane interaction assays.

### Formation of small-unilamellar vesicles (SUVs)

SUVs were prepared following the standard procedure as described in Sawa-Makarska et al., 2020. In brief, lipids were mixed in a glass vial and dried under an argon stream. The lipids were then dried further one hour under vacuum. After rehydration in 150 mM NaCl, 25mM Tris pH7.4 buffer the mixes were gently resuspended and sonicated for 2 min. The formed SUVs were then extruded first through a 0.4 μm, then a 0.1 μm membrane (Whatman, Nucleopore, UK) using the Mini Extruder from Avanti Polar Lipids Inc.. The final SUV suspension has a concentration of 1 mM or 2 mM.

### Microscopy-based protein-membrane interaction assays

For Figure 1E and 2C wild type or mutant versions of Faa1-EGFP were recruited to GFP-Trap_A beads beads (Chromotek). Assays were performed under equilibrium conditions with 1 mM of the prey liposomes in buffer (50 mM HEPES/Tris pH 7.5, 150 mM NaCl). The lipid mixtures were composed of 44 % POPC, 6 % POPS, 5.5 % POPE, 41.5 % liver PI and 2.5 % PI3P for Atg9 vesicle-like liposomes with PI3P, 44 % POPC, 6 % POPS, 5.5 % POPE and 44 % liver PI for Atg9 vesicle-like liposomes without PI3P, 63 % POPC, 6 % POPS, 5.5 % POPE and 21 % liver PI for Atg9 vesicle-like liposomes with 25 % PI, 83 % POPC, 6 % POPS, 5.5 % POPE and 5 % liver PI for Atg9 vesicle-like liposomes with 5 % PI, and 99.5 % POPC for POPC only liposomes. Additionally, they were labelled with 0.5 % lissamine-rhodamine-PE. Beads were imaged with a Zeiss LSM700 confocal microscope and a Plan-Apochromat 20x/0.8 objective. For quantification, lines were drawn across each bead in Fiji and the maximal values across the lines were taken. Values were averaged for each sample within each replicate, and then among replicates. The final values were then normalized to max. Beads from three independent experiments were quantified for the rhodamine signal. Significant differences were calculated with a Welch two-sided t-test.

For testing the recruitment of Faa1 to Atg9-PLs, Atg9-EGFP PLs were immobilized on GFP-TRAP_A beads (Chromotek). The assay was performed under equilibrium conditions with a prey concentration of 1 µM Faa1-mCherry. Imaging and analysis were carried out as stated above for the membrane-protein interaction assay.

### Liposome co-pelletation assay

5 µg of the Faa1 variants were incubated with 25 (Fig 1D) or 50 µL (Fig 2B) liposomes (compositions as stated in the microscopy-based membrane protein interaction assay) for 30 min at room temperature. The reaction was centrifuged at 68 000 rpm for 15 min at 22 °C (Beckman-Coulter Optima MAX, TLA100 rotor). The supernatant and pellet were loaded separately on an SDS/polyacrylamide gel in equal amounts. The quantification of the bands was performed with the Analyze gels tool of Fiji (Version 2.9.0/1.53t). The percentage of pelleted protein was calculated and the buffer control was subtracted. Significant differences were calculated with a Welch two-sided t-test.

### Faa1 activity assay

The activity of the Faa1 variants was determined by measuring the production of adenosine monophosphate (AMP) with an adapted protocol from Ford and Way, 2015. 5 µg of the Faa1 variants were incubated with 20 µL of 1 mM liposomes (44 % POPC, 6 % POPS, 6 % POPE, 41.5 % liver PI and 2.5 % PI3P for Atg9 vesicle-like liposomes with PI3P, 44 % POPC, 6 % POPS, 6 % POPE and 44 % liver PI for Atg9 vesicle-like liposomes without PI3P, and 100 % POPC for POPC only liposomes) for 30 min at room temperature. The reaction was mixed in a volume of 100 µL containing the Faa1-liposome mix, 8 mM MgCl2, 2 mM EDTA, 0.1 mM oleic acid, 0.2 mM NADH, 0.3 mM phosphoenolpyruvate, 0.1 mM bovine serum albumin, 2.5 mM ATP, 0.05 mM fructose 1,6-bisphosphate, 22.5 U myokinase from rabbit muscle (Sigma-Aldrich), 48 U pyruvate kinase from rabbit muscle (Sigma-Aldrich) and 24 U L-lactate dehydrogenase from rabbit muscle (Sigma-Aldrich) in buffer containing 150 mM NaCl and 25 mM Tris pH7.4. The reaction was induced by the addition of 0.5 mM coenzyme A or 0.5 mM AMP for the positive control. The absorbance at 340 nm was measured for 30 min at 30 °C with a Tecan SPARK Multimode Microplate Reader (Tecan Life Sciences). The reaction for FadD activity was performed with 5 µg FadD and without liposomes. For the experiment in Figure 3B Faa1-mCherry was used. For the other coupled enzymatic assays the EGFP-tagged variants were used. The measurements were normalized to the first timepoint before the initiation of the reaction.

### Yeast strains and media

Strains for *in vivo* analyses of Faa1 localization and function derive from genetic modification of *S. cerevisiae* w303. Knockouts and tags were obtained via homologous recombination of PCR products of interests as described previously (Kevin R. et al., 1997, Gietz et al., 1995). Briefly, log-phase cells were washed with lithium acetate (100 mM) and incubated in a mix containing lithium acetate (Sigma), PEG 2000 (Sigma), ssDNA and the PCR product of interest at 30 °C for 30’ and then shifted to 42 °C for 20’.

Strains were grown in synthetic complete medium (0.17 % (w/v) yeast nitrogen base without amino acids and ammonium sulfate (BD) supplemented with 0.5 % (w/v) ammonium sulfate (Sigma) 2 % (w/v) a-D-glucose monohydrate (Sigma) and complete supplement (CSM) drop out (Formedium)). Log-phase cells were harvested, washed 5 times and resuspended in SD-N medium (0.17 % (w/v) yeast nitrogen base without amino acids and ammonium sulfate (BD) and 2 % (w/v) a-D-glucose monohydrate (Sigma)). Based on the experiment, SD-N medium was supplemented with cerulenin (50 mg/mL; Merk) or glucose was reduced to 0.01 % (w/v). Rapamycin (400 ng/mL; Calbiochem) was added to synthetic complete medium.

### Cell viability

Cell suspension was mixed with 0.2 % (v/v) trypan blue (Invitrogen) and incubated for 5’. Cell viability was assessed by brightfield microscopy and expressed as percentage of blue-stained (dead) in 100 cells.

### Fluorescent microscopy

Cells were imaged at room temperature in a 96 well plate with glass bottom (Greiner Bio-One) using an inverted microscope Zeiss Axio Observer Z1 equipped with an EC Plan-Neofluar 100x/1.3 Oil M27 objective and an Orca Flash 4.0 LT+ camera. Images were deconvolved with Huygens Professional 23.04 (Scientific Volume Imaging) and analyzed using Fiji Version 2.9.0/1.53t.

### Western Blot and autophagic flux analysis

0.5 OD_600_ yeast cells were collected and lysed with 0.255 M NaOH. Proteins were precipitated with 100 % trichloroacetic acid and washed once with cold acetone. Protein pellets were resuspended in protein loading buffer (62.5 mM Tris-HCl pH 6.8, 10 % glycerol, 300 mM b-mercaptoethanol, 2 % SDS, 0.1 % Bromphenol-Blue sod salt) and analyzed by SDS-PAGE using mouse anti-GFP antibody (Max Perutz Labs, Monoclonal antibody facility) in 3 % (w/v) milk powder TBST-T (Sigma). Secondary antibody incubation was performed using Dylight 800 α-mouse (Rockland Immunochemicals). Detection and analysis were performed using the Li-COR Odyssey Infrared Imaging system (Biosciences) and the Image Studio (Version 2.1) software. Autophagic flux was assessed as the ratio between the free GFP and the total GFP signal (Free GFP and GFP Atg8 combined). Protein expression levels were normalized using Pgk1 as loading control (Invitrogen).

### CHARM-GUI HMMM Faa1-membrane model

Faa1-membrane interaction model was generated with CHARMM-GUI HMMM builder using following membrane composition: 44 % POPC, 41 % POPI, 6 % POPS, 6 % POPE, 3 % PI3P. Faa1 structural information was gathered from Alphafold2. Default ion concentration is 0.15 M.

**Fig. S1.**
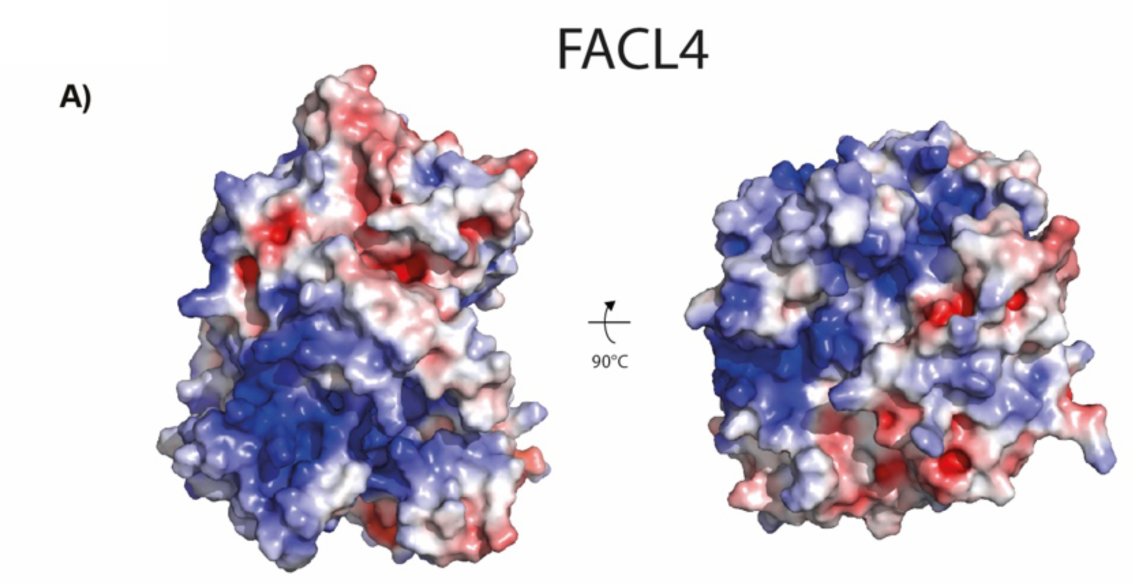
Alphafold model of human FACL4. (A) The distribution of the electrostatic potentials was calculated with the APBS method on the molecular surface of FACL4. Electro-positively and electronegatively charged areas are colored in blue and red, respectively. Neutral residues are depicted in white.

**Fig. S2.**
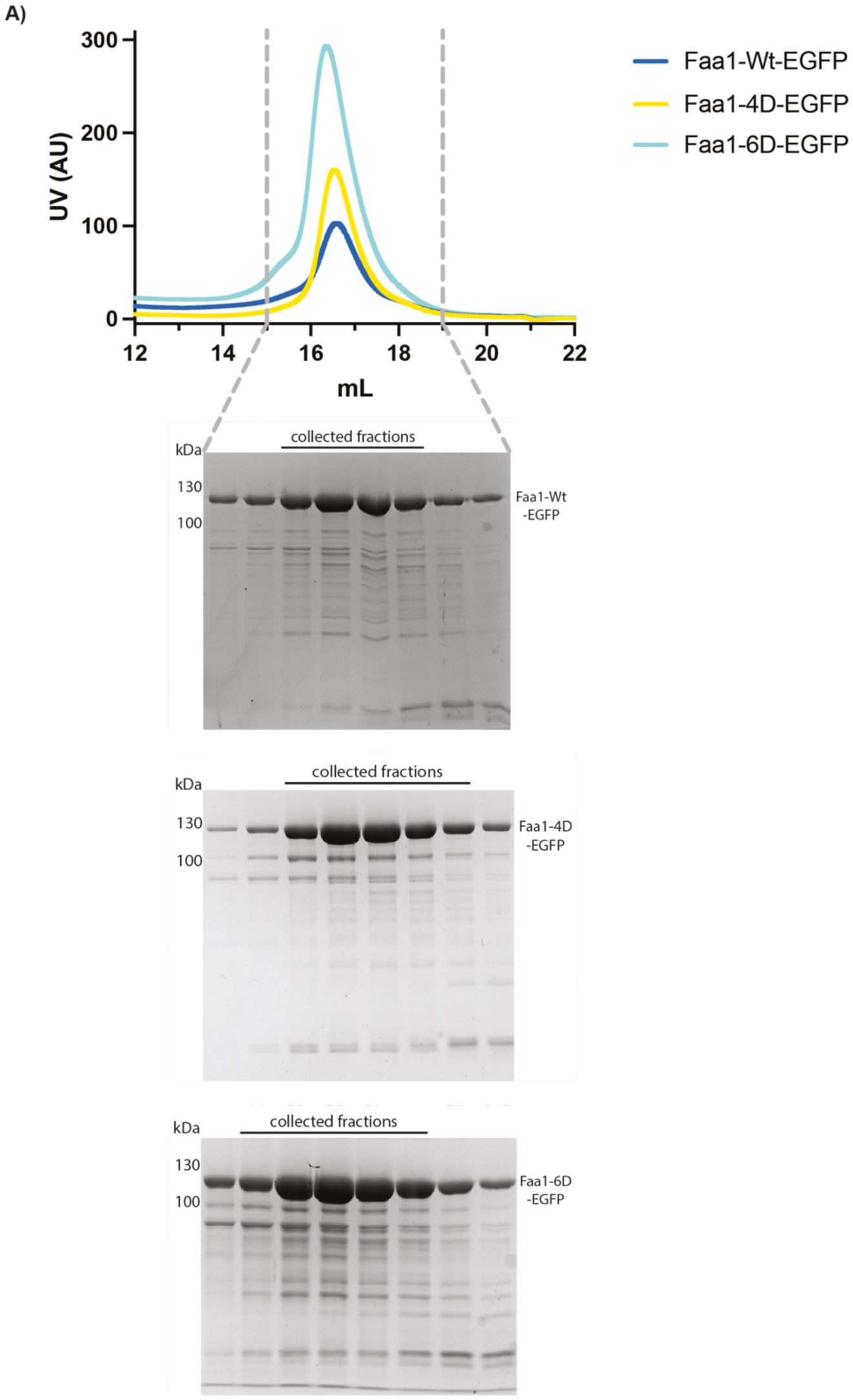
Size exclusion chromatography profile of Faa1 purifications. (A) The size exclusion chromatograms of Faa1-Wt-EGFP, Faa1-4D-EGFP and Faa1-6D-EGFP show that all three proteins purify identically indicating that the mutations do not cause misfolding or aggregation.

**Fig. S3.**
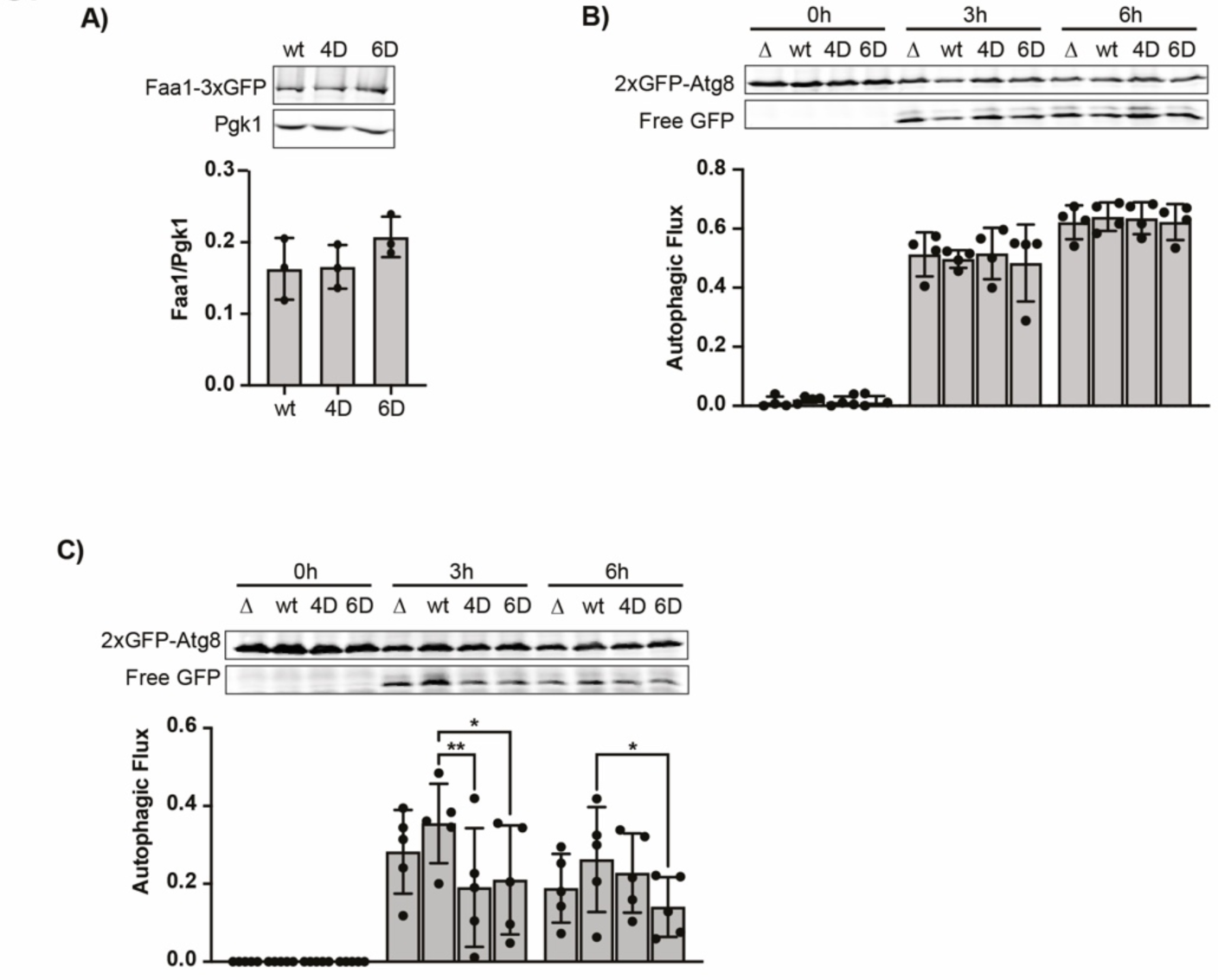
Mutation of positively charged residues in the plane area reduces Faa1 membrane localization and function in vivo. (A) Normalized protein levels of genomically tagged Faa1-wt, −4D and −6D by Western blot quantification. Data are means ± SD (n = 3). B) Autophagic flux of indicated strains expressing 2xGFP-ATG8 after 0, 3, 6h of starvation. Data are means ± SD (n = 4). (C) Autophagic flux of indicated strains expressing 2xGFP-ATG8 after 0, 3, 6h of starvation and glucose depletion (0.01% w/v). Data are means ± SD (n = 5). One-way ANOVA: *<0.05, **<0.01, ***<0.001

